# Atherosis of trophoblast type: A specific form of decidual vasculopathy distinct from atherosis of macrophage type

**DOI:** 10.1101/2021.07.02.450868

**Authors:** Peilin Zhang

## Abstract

**Background:** There are three types of decidual vasculopathy, namely, acute atherosis, fibrinoid medial necrosis and mural arterial hyerptrophy. Persistence of vascular trophoblasts is also known to be related to maternal vascular malperfusion, but detailed study is lacking.

**Material and methods:** A total 1017 placentas from 2021 were collected with clinical, neonatal and placental information, and routine placental pathology examination was performed. Decidual vasculopathy was classified based on the new classification scheme including atherosis of macrophage type atherosis of trophoblast type, fibrinoid medial necrosis, mural arterial hypertrophy and mixed type vasculopathy. The significance of these morphologic changes were examined based on the clinical, neonatal and placental pathology features.

**Results:** Decidual vasculopathy is classified as classic type, mural hypertrophy and mixed type. Classic type vasculopathy is further separated as atherosis and fibrinoid medial necrosis. Atherosis is defined as atherosis of macrophage type and atherosis of trophoblast type. Each category of decidual vasculopathy was evaluated in association with maternal, neonatal and placental pathologic findings. Atherosis of macrophage type and mixed type vasculopathy showed statistically significant association with preeclampsia/pregnancy induced hypertension, low birth weight and low placental weight. Atherosis of trophoblast type was associated with lower placental weight but not with specific clinical features. There is no neonatal sex dimorphism in decidual vasculopathy.

**Conclusion:** Atherosis of trophoblast type is a distinct pathologic feature in late pregnancy, and it is associated with lower placental weight. New classification of decidual vasculopathy helps with better stratification and categorization of placental maternal vascular abnormalities.

## Introduction

Maternal vascular transformation is a hallmark of human pregnancy and represents critical steps towards normal embryonic and placental development [1]. Abnormal maternal vascular transformation is invariably associated with adverse pregnancy complications. In current guideline of placental examination, abnormal maternal vascular transformation is grouped under the heading of maternal vascular malperfusion characterized by a constellation of morphologic changes in placental tissue after delivery [2]. There are three distinct types of maternal vascular changes recognized in the current guideline; namely acute atherosis, fibrinoid medial necrosis and mural arterial hypertrophy [2–4]. Persistence of intramural endovascular trophoblasts in third trimester is also recognized as a distinct form of decidual arteriopathy, although the pathogenic mechanism of all these morphologic changes is unclear [2, 5]. Acute atherosis, first described by Zeek, is characterized by the presence of foamy macrophages in the arterial wall with or without fibrinoid medial necrosis, similar to those in atherosclerosis in adult [6]. Acute atherosis is more commonly identified in the fetal membranes (decidua capsularis) and the foamy macrophages within the arterial walls are thought to be related to oxidative stress [6–9]. Acute atherosis is associated with preeclampsia and hypertensive disorders of pregnancy [1, 6, 10]. We have previously shown that a subset of patients with acute atherosis demonstrated the presence of foamy trophoblasts within the wall or lumen of spiral arteries (intramural or endovascular), and these foamy trophoblasts express surface CD56, a natural killer cell marker [11–14]. Recognition of foamy trophoblasts in acute atherosis raises new questions about the pathogenesis. In addition, persistence of intramural or endovascular trophoblasts in the “remodeled spiral arteries” at late pregnancy is far more common, and when present in the superficial myometrium, it was named as “physiologic changes” with implication of normal pregnancy findings [5, 15, 16]. Given the controversial nature of these morphologic changes historically, it is necessary to revisit the issue in light of endovascular trophoblastic expression of CD56, to separate the atherosis of macrophage type (Zeek’s atherosis) from atherosis of trophoblast type, and to properly define the term of atherosis of trophoblast type. Separation of atherosis of macrophage type from trophoblast type raises new questions of pathogenesis and provides new direction of research in maternal risk assessment of cardiovascular disease later in life.

## Material and methods

The study is exempt from Institutional Review Board (IRB) approval according to section 46.101(b) of 45CFR 46 which states that research involving the study of existing pathological and diagnostic specimens in such a manner that subjects cannot be identified is exempt from the Department of Health and Human Services Protection of Human Research Subjects. Placental tissues submitted for pathology examination for a variety of clinical indications in 2021 were included in the study using the routine Hematoxylin & Eosin (H&E) stain. Paraffin-embedded tissues from routine surgical pathology specimens and routine H&E stained pathology slides were examined by light microscopy using the Amsterdam criteria for placental examination [1, 2]. Immunohistochemical staining for CD56 and CD68 was used to highlight the specific cell types when needed. Immunohistochemical staining procedures were performed on paraffin-embedded tissues using Leica Biosystems Bond III automated immunostaining system following the manufacturer’s instructions [11]. CD56 and CD68 monoclonal antibodies were purchased for clinical applications from Dako Agilent under the catalogue number M730401-2 and M087601-2.

A total 1017 placentas with corresponding neonatal data from Jan. 2021 to Jun 2021 were collected from the medical records including 96 normal pregnancies. The placental pathology data was entered into Excel spreadsheet at the time of pathology examination, and the neonatal sex, birth weight, birth length and head circumference were subsequently retrieved from the medical records. Twin pregnancies with identical sex were included in the study, and the twin pregnancies with different sex were excluded.

Separately, a database of pre-eclampsia/pregnancy induced hypertension was established between 2017 to 2020 consisting of placental pathologic features and clinical features. The data include 619 placentas of late onset preeclampsia and 133 normal controls. A portion of these data was previously published [11]. Atherosis within this data set was collectively named as classic type without further separation of atherosis of macrophage type and atherosis of trophoblast type. Early onset preeclampsia was excluded from current study due to the confounding factor of gestational age affecting the placental and neonatal weight.

Statistical analysis was performed by using various programs of R-Package including baseline characteristic table and multi-variant ANOVA tests (http://statistics4everyone.blogspot.com/2018/01/fathers-data-visualization.html).

## Results

### 1. Atherosis of macrophage type and atherosis of trophoblast type

In typical acute atherosis, foamy macrophages can be identified within the arterial walls and these macrophages can be highlighted by immunohistochemical staining for CD68, a macrophage marker (Figure 1, top panel). Acute atherosis was initially described by Zeek [6], and the terminology remains essentially unchanged to date with little controversy. In a subset of acute atherosis, the foamy cells were not proven to be macrophages but endovascular trophoblasts with unique CD56 expression (Figure 1, bottom panel) [11]. CD56 expression is restricted on the endovascular trophoblasts only, in addition to decidual natural killers cells (NK), thus separating the population of extravillous trophoblasts from that of endovascular trophoblasts [11]. Atherosis of macrophage type shares significant morphologic similarities to atherosis of trophoblast type, and both can show foamy cell changes in the background of eosinophilic fibrinoid arterial media (fibrinoid medial necrosis). The morphologic differences can be appreciated with slightly basophilic cytoplasm of intramural or endovascular trophoblasts under light microscopy whereas the clear foamy cytoplasm of macrophages is normally identified. In Zeek’s initial description of atherosis of macrophage type, anatomic location of membrane rolls was emphasized [6], consistent with the lack of trophoblast invasion and trophoblast-directed spiral artery remodeling in the decidual capsularis.

**Figure 1.**
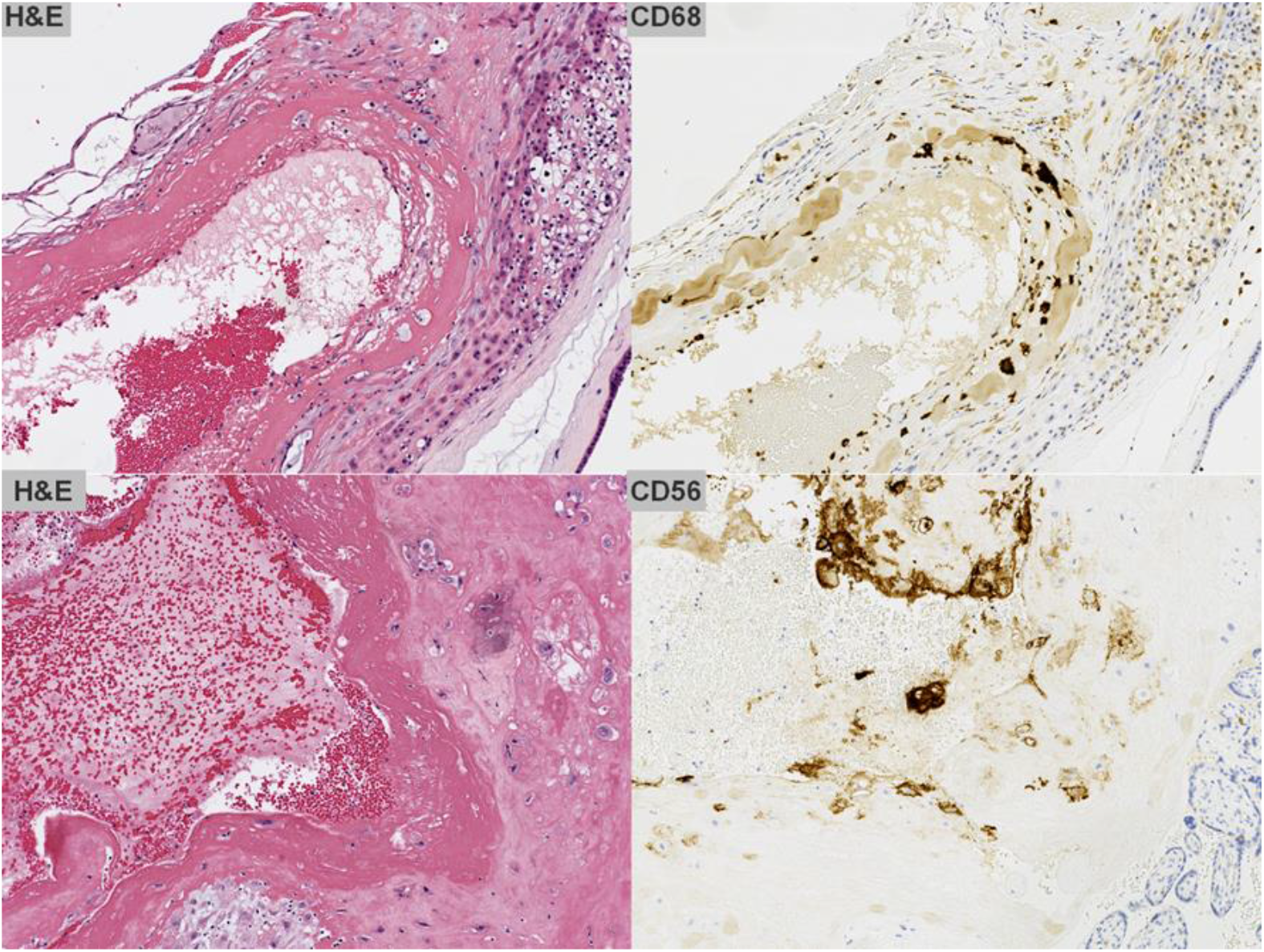
Atherosis of trophoblast type and atherosis of macrophage type. Typical atherosis of macrophage type (top panel) with CD68 expression and atherosis of trophoblast type with CD56 expression by immunohistochemical staining. Magnification 200X.

### 2 New classification of decidual vasculopathy

Based on the separation of atherosis of macrophage type from trophoblast type, the decidual vasculopathy can be classified as classic type, mixed type and mural arterial hypertrophy as in Table 1. Atherosis and fibrinoid medial necrosis co-exist commonly, and pure atherosis without fibrinoid medial necrosis is rare. Both atherosis and fibrinoid medial necrosis are historic terms, and so named as “classic type”. The term of atherosis of trophoblast type still applies even when the background of fibrinoid medial necrosis in late pregnancy is absent, given the fact that the pathogenic mechanism is the persistence of vascular trophoblasts in third trimester. Mural arterial hypertrophy is a separate identity and it is independent of trophoblast invasion, and typically occurs in the small caliber arteries in either decidua basalis or capsularis. Mural arterial hypertrophy may relate to hormonal influence (progesterone) as similar changes of small arteries can be seen in secretory endometrium without pregnancy [17]. It is worth noting that morphologic changes of mural hypertrophy in superficial myometrium were used as evidence of “failed trophoblast invasion” in study of preeclampsia [15, 16].

**Table 1:**
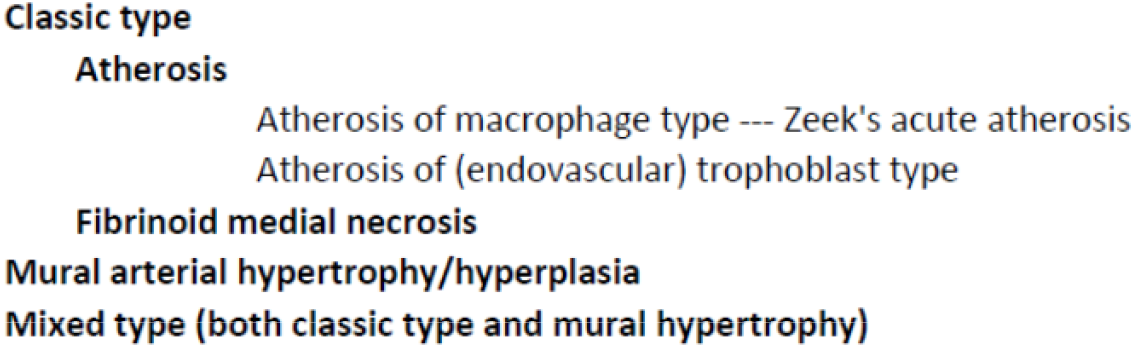
Types of decidual vasculopathy. Classification of decidual vasculopathy.

Mixed type vasculopathy is defined as the presence of both atherosis and mural arterial hypertrophy in the same or different anatomic locations such as decidual basalis and/or decidual capsularis. The morphologic features of atherosis and mural arterial hypertrophy are different with different implications of pathogenesis, and the combination of both morphologic features (Figure 2 and Figure 3) in the same case represents a distinct risk for both the mothers and fetus. In our data, mixed type vasculopathy is associated with most significant changes in placental weight, neonatal weight and growth as well as placental pathology (later section).

**Figure 2.**
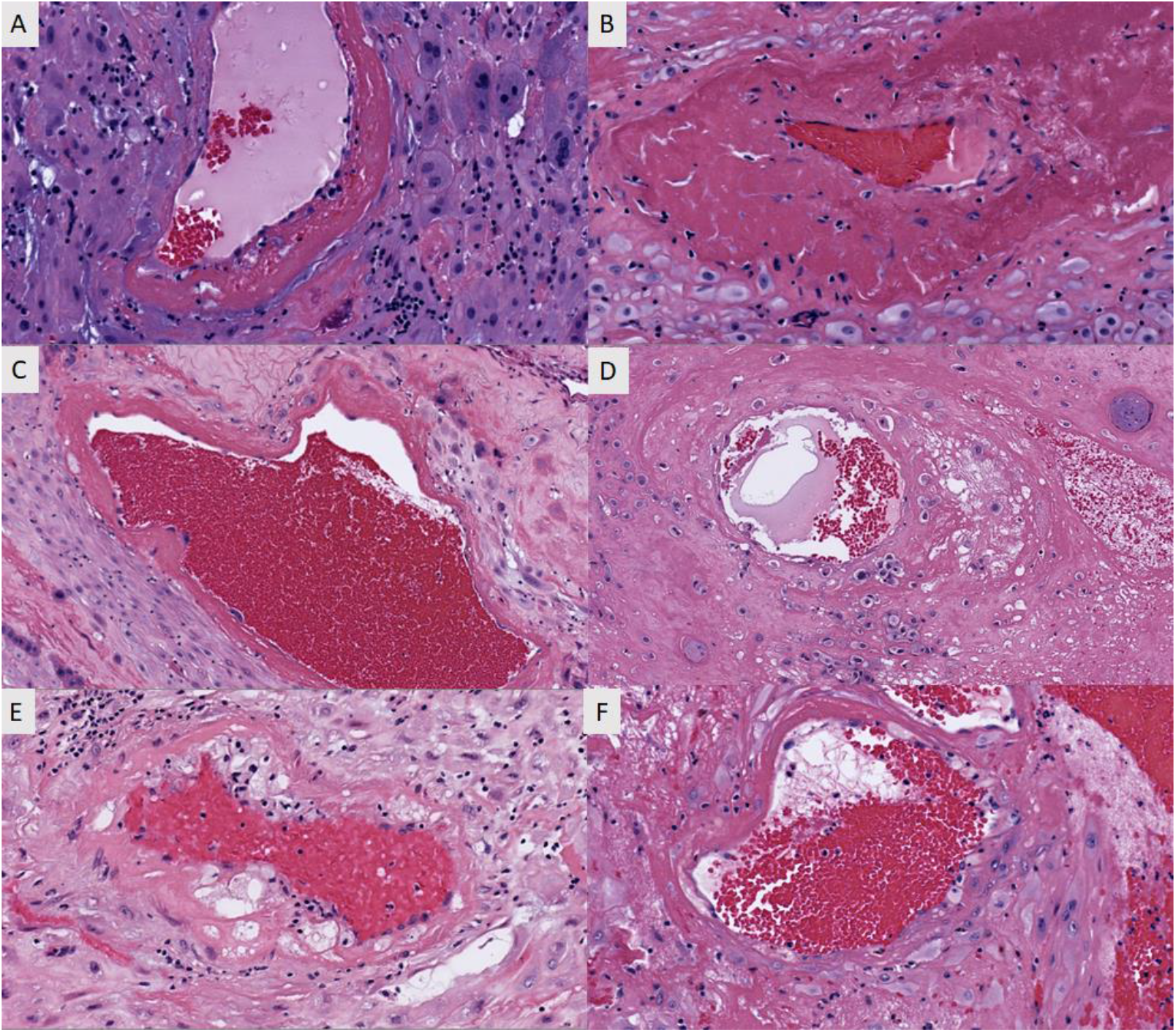
Morphologic variant of atherosis of trophoblast type. Multiple morphologic variants of atherosis of trophoblast type with various degrees of fibrinoid medial necrosis associated with various numbers of intramural or endovascular trophoblasts (A-F). Magnification 200X.

**Figure 3.**
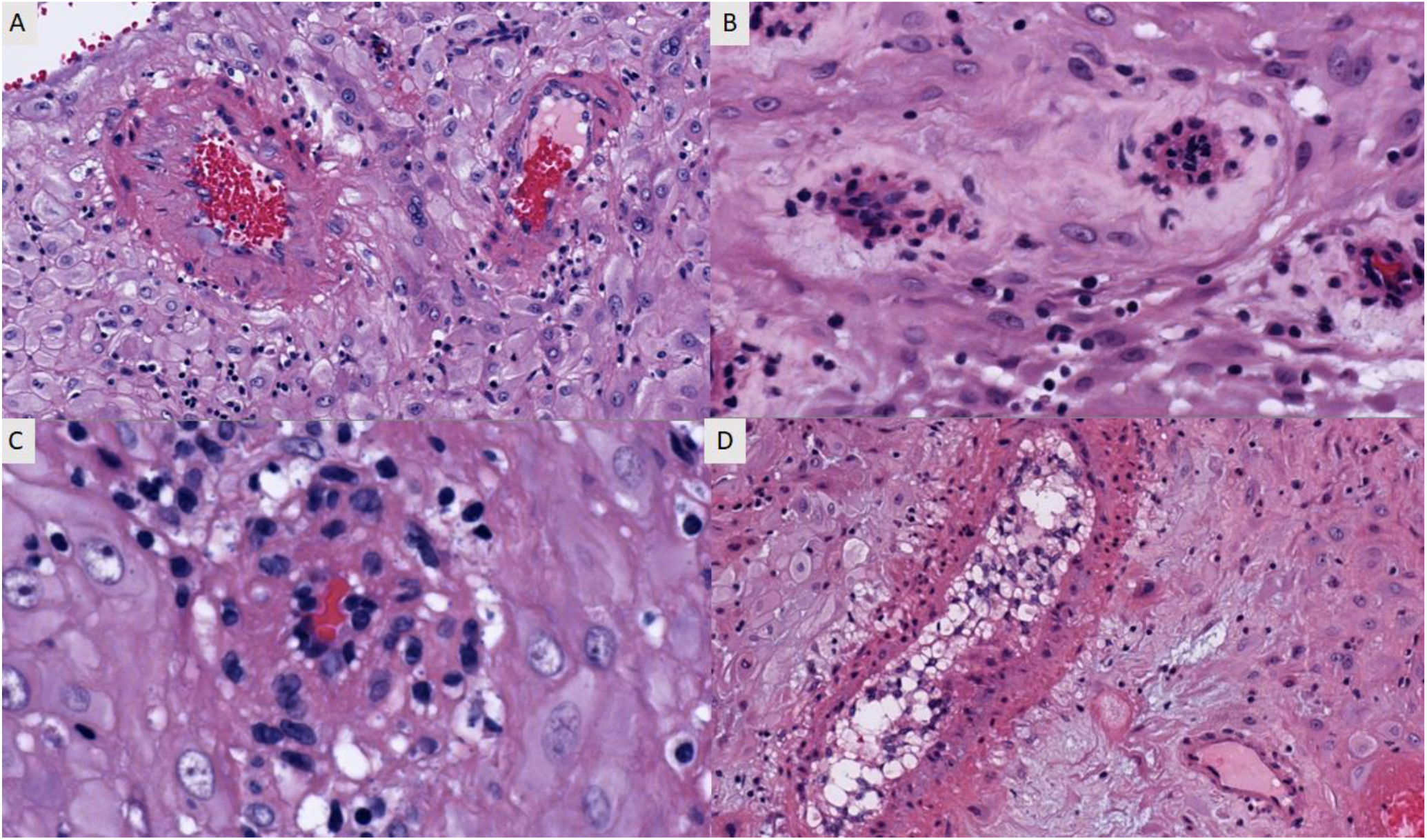
Morphologic variant of mural arterial hypertrophy. Multiple morphologic variants of mural arterial hypertrophy (A-D). Magnification 200X.

This new classification of decidual vasculopathy incorporated some new evidence of pathogenesis of maternal vascular transformation from early to late pregnancy and provided new insight in understanding of the interaction between the trophoblasts, endothelial cells and smooth muscle cells and potential risk of maternal cardiovascular disease in later life after pregnancy.

### 3, Morphologic variants of atherosis of trophoblast type

Practically, there are morphologic variants of atherosis of trophoblast type. The examples of these vascular changes are shown in Figure 2. In membrane rolls the decidual vessels are not remodeled by endovascular trophoblasts and the vascular changes including hypertrophic/hyperplastic and/or atherosis/fibrinoid medial necrosis are not directly related to trophoblasts, therefore atherosis of macrophages and mural hypertrophy/hyperplasia are present exclusively. In decidua basalis where the extensive spiral artery remodeling occurred at early implantation three types of vascular changes, both atherosis of trophoblasts and atherosis of macrophages, as well as mural hypertrophy can be identified. In our experience, atherosis of trophoblasts type is far more common than that of macrophage type in hypertensive or nonhypertensive disorders of pregnancy, suggesting significantly different pathogenic mechanisms of both types of atherosis. Unlike atherosis of macrophage type, atherosis of trophoblast type does not seem to play significant roles in pathogenesis of preeclampsia or hypertensive disorders of pregnancy.

### 4, Decidual vasculopathy and associated clinical and pathologic features

A total 1017 placentas with clinical, pathologic and neonatal data were collected in 2021 chronologically in light of new classification of decidual vasculopathy (Table 2). A total 339 of the 1017 placentas show features of various decidual vasculopathy. Atherosis of macrophage type (p=0.009) and mixed type vasculopathy (p=0.006) showed statistically significant association with preeclampsia/PIH. All types of decidual vasculopathy were associated with decreased placental weight, neonatal weight and IUGR (p=0.001). Most of the statistically significant associations between the decidual vasculopathy and clinical and placental pathologic features were previously known, and our current data is consistent with previous knowledge. There was statistically significant association of various types of decidual vasculopathy with BMI. There were statistically significant association between the specific vasculopathy type and placental weight at delivery. It is worth noting that mixed type vasculopathy is the most significant placental lesion and it is associated with lowest placental weight, neonatal weight, neonatal birth length and most significant placental pathologies.

**Table 2.** Decidual vasculopathy types and associated clinical and pathologic features. Abbreviation: GBS – Group B streptococcus; IUGR - Intrauterine growth restriction; IUFD – Intrauterine fetal demise; MIR – Maternal inflammatory response, FIR – Fetal inflammatory response; MFI – Maternal floor infarction /massive peri-villous fibrin deposit. Statistical analysis was performed by using baseline characteristic table in R-package and multi-variant ANOVA tests. P<0.05 is statistically significant.

| Vasculopathy | Classic M<br>(N=10) | Classic T<br>(N=218) | Mixed type<br>(N=45) | Mural hypertrophy<br>(N=66) | No vasculopathy<br>(N=582) | Normal controls<br>(N=96) | p value |
| --- | --- | --- | --- | --- | --- | --- | --- |
| Neonatal sex |  |  |  |  |  |  | 0.268 |
| - Female | 3 (30.000%) | 108 (49.541%) | **22 (48.889%)** | 41 (62.121%) | 285 (48.969%) | 52 (54.167%) |  |
| - Male | 7 (70.000%) | 110 (50.459%) | **23 (51.111%)** | 25 (37.879%) | 297 (51.031%) | 44 (45.833%) |  |
| Neonatal weight (gram) | 2955.000 [2475.000;3580.000] | 3105.000 [2730.000;3530.000] | **2795.000 [2505.000;3290.000]** | 3270.000 [2850.000;3470.000] | 3270.000 [2830.000;3610.000] | 3160.000 [2700.000;3550.000] | **0.001** |
| Neonatal length (cm) | 50.000 [49.500;52.000] | 50.000 [48.000;51.000] | **49.000 [47.000;50.500]** | 50.000 [48.500;51.000] | 50.500 [48.000;52.000] | 50.000 [48.000;51.500] | **0.001** |
| Neonatal head circumference (cm) | 33.000 [32.500;33.500] | 34.000 [32.500;34.750] | **33.500 [32.000;34.000]** | 34.000 [32.500;35.000] | 34.000 [33.000;35.000] | 34.000 [33.000;35.000] | **0.018** |
| Umbilical cord length (cm) | 39.500 [33.000;42.000] | 35.000 [27.000;43.000] | **34.000 [29.500;39.000]** | 33.500 [30.000;42.000] | 36.000 [27.000;43.000] | 36.000 [29.000;43.000] | 0.698 |
| Cord coiling per 10 cm | 3.000 [3.000;4.000] | 3.000 [3.000;5.000] | **3.000 [3.000;5.000]** | 4.000 [3.000;5.000] | 4.000 [3.000;5.000] | 3.000 [3.000;5.000] | 0.827 |
| **Clinical features** |  |  |  |  |  |  |  |
| GBS status | 4 (40.000%) | 60 (33.898%) | **8 (24.242%)** | 21 (40.385%) | 158 (32.917%) | 0 ( 0.0%) |  |
| SARS-CoV2 status | 1 (10.000%) | 13 (5.963%) | **1 (2.222%)** | 3 (4.545%) | 30 (5.155%) | 7 (7.292%) | 0.810 |
| Maternal age (year) | 31.500 [26.000;34.000] | 32.000 [28.000;36.000] | **31.000 [27.000;33.000]** | 31.000 [27.000;35.000] | 32.000 [27.000;36.000] | 31.000 [26.500;35.000] | 0.627 |
| Delivery mode |  |  |  |  |  |  | 0.371 |
| - Cesarean | 5 (50.000%) | 75 (34.404%) | **16 (35.556%)** | 29 (43.939%) | 201 (34.536%) | 27 (28.125%) |  |
| - Vaginal | 5 (50.000%) | 143 (65.596%) | **29 (64.444%)** | 37 (56.061%) | 381 (65.464%) | 69 (71.875%) |  |
| Preeclampsia /PIH | 5 (50.000%) | 42 (19.266%) | **15 (33.333%)** | 14 (21.212%) | 85 (14.605%) | 3 (3.125%) | **0.000** |
| Gestational age (weeks) | 38.000 [37.000;40.000] | 39.000 [37.000;40.000] | **39.000 [37.000;40.000]** | 39.000 [38.000;40.000] | 39.000 [38.000;40.000] | 39.500 [39.000;40.000] | **0.012** |
| GDM2 | 2 (20.000%) | 31 (14.220%) | **8 (17.778%)** | 13 (19.697%) | 75 (12.887%) | 1 (1.042%) | **0.005** |
| Category 2 fetal heart tracing | 0 ( 0.0%) | 39 (17.890%) | **5 (11.111%)** | 10 (15.152%) | 111 (19.072%) | 0 ( 0.0%) | **0.000** |
| IUFD | 1 (10.000%) | 0 ( 0.0%) | **2 (4.444%)** | 0 ( 0.0%) | 11 (1.890%) | 0 ( 0.0%) | **0.010** |
| Oligohydramnios | 0 ( 0.0%) | 3 (1.376%) | **2 (4.444%)** | 1 (1.515%) | 16 (2.749%) | 0 ( 0.0%) | 0.396 |
| **Placental pathology** |  |  |  |  |  |  |  |
| Placenta weight (gram) | 460.000 [360.000;527.000] | 442.500 [375.000;515.000] | **416.000 [375.000;466.000]** | 479.000 [409.000;552.000] | 465.000 [390.000;544.000] | 488.500 [414.000;538.500] | **0.001** |
| **Maternal vascular malperfusion** |  |  |  |  |  |  |  |
| Infarcts | 2 (20.000%) | 19 (8.716%) | **6 (13.333%)** | 2 (3.030%) | 37 (6.357%) | 5 (5.208%) | 0.117 |
| Thrombosis | 4 (40.000%) | 81 (37.156%) | **13 (28.889%)** | 9 (13.636%) | 113 (19.449%) | 18 (18.750%) | **0.000** |
| Abruption | 1 (10.000%) | 3 (1.376%) | **2 (4.444%)** | 0 ( 0.0%) | 8 (1.375%) | 2 (2.083%) | 0.135 |
| IUGR | 1 (10.000%) | 15 (6.912%) | **7 (15.556%)** | 4 (6.061%) | 27 (4.639%) | 0 ( 0.0%) | **0.005** |
| **Fetal vascular malperfusion** |  |  |  |  |  |  |  |
| Avascular villi | 0 ( 0.0%) | 23 (10.550%) | **5 (11.111%)** | 6 (9.091%) | 83 (14.261%) | 12 (12.500%) | 0.476 |
| **Inflammatory/infectious** |  |  |  |  |  |  |  |
| MIR (acute chorioamnionitis) | 3 (30.000%) | 56 (25.688%) | **7 (15.556%)** | 9 (13.636%) | 184 (31.615%) | 35 (36.458%) | **0.004** |
| MIR (chronic deciduitis) | 1 (10.000%) | 42 (19.266%) | **9 (20.000%)** | 12 (18.182%) | 120 (20.619%) | 44 (45.833%) | **0.000** |
| FIR (acute funisitis/fetal vasculitis) | 1 (10.000%) | 22 (10.092%) | **3 (6.667%)** | 1 (1.515%) | 81 (13.918%) | 12 (12.500%) | 0.051 |
| Chronic villitis | 1 (10.000%) | 31 (14.220%) | **5 (11.111%)** | 11 (16.667%) | 112 (19.244%) | 30 (31.250%) | **0.009** |
| Other pathology |  |  |  |  |  |  |  |
| Meconium stain | 0 ( 0.0%) | 53 (24.424%) | **4 (8.889%)** | 19 (28.788%) | 167 (28.694%) | 26 (27.083%) | **0.024** |
| MFI/MPFD | 0 ( 0.0%) | 5 (2.294%) | **1 (2.222%)** | 2 (3.030%) | 19 (3.265%) | 3 (3.158%) | 0.968 |
| **Lab and other tests** |  |  |  |  |  |  |  |
| Total WBC count | 9.300 [8.000;10.100] | 9.700 [8.100;11.500] | 9.100 [7.750;11.900] | 9.300 [7.800;11.050] | 9.700 [8.200;12.300] | 10.000 [8.700;12.800] | 0.274 |
| Neutrophil differentials | 72.750 ± 4.459 | 71.731 ± 7.136 | 71.381 ± 8.521 | 71.025 ± 6.190 | 72.493 ± 7.460 | 74.633 ± 7.553 | 0.131 |
| Body temperature (°C) | 36.850 [36.500;37.000] | 36.700 [36.500;36.900] | 36.800 [36.600;37.000] | 36.700 [36.500;37.000] | 36.700 [36.500;36.900] | 36.800 [36.500;37.100] | 0.184 |
| BMI | 32.876 [31.030;35.929] | 30.105 [27.125;33.871] | 30.852 [27.467;37.863] | 33.775 [30.040;36.946] | 30.731 [27.493;35.177] | - | **0.004** |
| Blood pressure (Systolic) | 129.500 [125.000;145.000] | 128.000 [118.000;137.000] | 128.000 [122.000;141.000] | 129.500 [118.000;137.000] | 126.500 [118.000;135.000] | - | 0.262 |
| Blood pressure (Diastolic) | 79.500 [76.000;83.000] | 79.000 [71.000;85.000] | 79.000 [73.000;88.000] | 76.000 [71.000;87.000] | 78.000 [70.000;84.000] | - | 0.546 |

### 5. Neonatal sex and decidual vasculopathy

Atherosis of trophoblast type with persistence of vascular trophoblasts in late pregnancy leads to questions of failed trophoblast cell death, or failed endothelial cell/smooth muscle cell regeneration or both. We explored the potential association between the neonatal sex and atherosis of trophoblast type, as vascular trophoblasts are of fetal origin in maternal circulation. Trophoblast invasion into spiral artery and persistence of endovascular trophoblasts within maternal spiral artery may relate to sex dimorphism. As shown in Table 3, there is no statistically significant difference in frequencies of all types of decidual vasculopathy between male and female neonates. There is no difference in placental pathology and lab tests except for neonatal birth weight, birth length and birth head circumference between male and female sex. There is statistically higher incidence of Cesarean section for male neonate than female neonates.

**Table 3.** Neonatal sex and associated clinical and pathologic features. Abbreviation: GBS – Group B streptococcus; IUGR - Intrauterine growth restriction; IUFD – Intrauterine fetal demise; MIR – Maternal inflammatory response, FIR – Fetal inflammatory response; MFI – Maternal floor infarction /massive peri-villous fibrin deposit. Statistical analysis was performed by using baseline characteristic table in R-package and multi-variant ANOVA tests. P<0.05 is statistically significant.

| Neonatal sex | Female | Male | p value |
| --- | --- | --- | --- |
|  | (N=511) | (N=506) |  |
| Neonatal weight (gram) | 3140.000 [2745.000;3530.000] | 3260.000 [2815.000;3580.000] | 0.045 |
| Neonatal length (cm) | 50.000 [48.000;51.500] | 50.500 [48.000;52.000] | 0.001 |
| Neonatal head circumference (cm) | 33.800 [32.500;34.500] | 34.000 [33.000;35.000] | 0.000 |
| Umbilical cord length (cm) | 35.000 [27.000;42.000] | 36.000 [28.000;43.000] | 0.098 |
| Cord coiling per 10 cm | 4.000 [3.000;5.000] | 4.000 [3.000;5.000] | 0.206 |
| **Clinical features** |  |  |  |
| GBS status | 135 (35.156%) | 116 (31.522%) | 0.327 |
| SARS-CoV2 status | 33 (6.458%) | 22 (4.348%) | 0.177 |
| Maternal age (year) | 32.000 [27.000;35.000] | 32.000 [27.000;36.000] | 0.499 |
| Gestational age (week) | 39.000 [38.000;40.000] | 39.000 [38.000;40.000] | 0.750 |
| Delivery |  |  | 0.019 |
| - Cesarean | 159 (31.115%) | 194 (38.340%) |  |
| - Vaginal | 352 (68.885%) | 312 (61.660%) |  |
| Preeclampsia/PIH | 92 (18.004%) | 72 (14.229%) | 0.121 |
| GDM2 | 65 (12.720%) | 65 (12.846%) | 1.000 |
| Category 2 fetal heart tracing | 77 (15.068%) | 88 (17.391%) | 0.358 |
| IUGR | 30 (5.882%) | 24 (4.743%) | 0.503 |
| IUFD | 6 (1.174%) | 8 (1.581%) | 0.774 |
| Oligohydramnios | 12 (2.348%) | 10 (1.976%) | 0.848 |
| **Placental pathology** |  |  |  |
| Maternal vascular malperfusion |  |  |  |
| Decidual vasculopathy |  |  | 0.268 |
| Classic (Macrophage) | 4 (0.860%) | 5 (1.080%) | 0.995 |
| Classic (Trophoblast) | 108 (23.529%) | 111 (24.026%) | 0.921 |
| Mixed type | 21 (4.575%) | 23 (4.978%) | 0.895 |
| Mural hypertrophy | 42 (9.150%) | 26 (5.628%) | 0.055 |
| Placental weight (gram) | 463.000 [386.000;538.500] | 460.500 [392.000;531.000] | 0.994 |
| Infarcts | 35 (6.849%) | 36 (7.115%) | 0.966 |
| Thrombosis | 121 (23.679%) | 117 (23.168%) | 0.906 |
| Abruption | 6 (1.174%) | 10 (1.976%) | 0.438 |
| Fetal vascular malperfusion |  |  |  |
| Avascular villi | 69 (13.503%) | 60 (11.858%) | 0.488 |
| Inflammatory/infectious |  |  |  |
| MIR (acute chorioamnionitis) | 139 (27.202%) | 155 (30.632%) | 0.255 |
| MIR (chronic deciduitis) | 113 (22.114%) | 115 (22.727%) | 0.873 |
| FIR (acute funisitis/vasculitis) | 55 (10.763%) | 65 (12.846%) | 0.351 |
| Chronic villitis | 99 (19.374%) | 91 (17.984%) | 0.626 |
| **Other placental pathology** |  |  |  |
| Meconium stain | 147 (28.824%) | 122 (24.111%) | 0.103 |
| MFI/MPFD | 15 (2.941%) | 15 (2.964%) | 1.000 |
| **Lab tests and others** |  |  |  |
| Total WBC counts | 9.600 [8.100;11.700] | 9.900 [8.100;12.300] | 0.121 |
| Neutrophil differential (%) | 72.116 ± 7.042 | 72.422 ± 7.661 | 0.525 |
| Lymphocyte differential (%) | 17.400 [13.150;21.650] | 17.300 [13.000;21.200] | 0.819 |
| Body temperature (°C) | 36.700 [36.500;36.900] | 36.700 [36.500;36.900] | 0.751 |
| BMI | 30.825 [27.453;34.874] | 30.828 [27.590;35.855] | 0.431 |
| Blood pressure (Systolic) | 126.000 [118.000;137.000] | 127.500 [119.000;136.000] | 0.391 |
| Blood pressure (Diastolic) | 77.500 [70.000;84.000] | 78.500 [72.000;85.000] | 0.341 |

### 6. Association of preeclampsia/PIH with decidual vasculopathy

Atherosis of macrophage type is known to be associated with preeclampsia/PIH [6]. We examine our current dataset to see if there is an association of atherosis of trophoblast type with preeclampsia/PIH (Table 4). There is statistically significant association between atherosis of macrophage type, mixed type vasculopathy and preeclampsia (p=0.009, and p=0.006 respectively). There is no association between atherosis of trophoblast type, mural arterial hypertrophy and preeclampsia (p=0.512 and p=0.593 respectively). There are many positive and negative associations between clinical and pathologic features with preeclampsia, and most of these associations are previously known (Table 4).

**Table 4.** Preeclampsia/PIH and associated clinical and pathologic features. Abbreviation: PIH – Pregnancy induced hypertension; IUGR - Intrauterine growth restriction; IUFD – Intrauterine fetal demise; GBS – Group B streptococcus; MIR – Maternal inflammatory response, FIR – Fetal inflammatory response; MFI – Maternal floor infarction /massive peri-villous fibrin deposit. Statistical analysis was performed by using baseline characteristic table in R-package and multi-variant ANOVA tests. P<0.05 is statistically significant.

| **Preeclampsia/PIH** | **Absent**<br>(N=853) | **Present**<br>(N=164) | **p value** |
| --- | --- | --- | --- |
| Neonatal sex |  |  | 0.121 |
| - Female | 419 (49.121%) | 92 (56.098%) |  |
| - Male | 434 (50.879%) | 72 (43.902%) |  |
| Neonatal weight (grams) | 3240.000 [2810.000;3585.000] | 2940.000 [2605.000;3440.000] | 0.000 |
| Neonatal length (cm) | 50.000 [48.000;52.000] | 50.000 [48.000;51.000] | 0.013 |
| Neonatal head circumference (cm) | 34.000 [33.000;35.000] | 33.500 [32.000;34.500] | 0.011 |
| Umbilical cord length (cm) | 35.000 [28.000;42.000] | 37.000 [29.500;43.500] | 0.108 |
| Cord coiling per 10 cm | 4.000 [3.000;5.000] | 3.000 [3.000;5.000] | 0.276 |
| **Clinical features** |  |  |  |
| GBS status | 202 (31.861%) | 49 (41.525%) | 0.053 |
| SARS-CoV2 status | 49 (5.744%) | 6 (3.659%) | 0.372 |
| Maternal age (year) | 32.000 [27.000;35.000] | 32.000 [28.000;37.000] | 0.190 |
| Gestational age (week) | 39.000 [38.000;40.000] | 38.000 [37.000;39.500] | 0.000 |
| Delivery |  |  | 0.522 |
| - Cesarean | 292 (34.232%) | 61 (37.195%) |  |
| - Vaginal | 561 (65.768%) | 103 (62.805%) |  |
| GDM2 | 115 (13.482%) | 15 (9.146%) | 0.163 |
| Category 2 fetal heart tracing | 159 (18.640%) | 6 (3.659%) | 0.000 |
| IUGR | 46 (5.393%) | 8 (4.908%) | 0.950 |
| IUFD | 12 (1.407%) | 2 (1.220%) | 1.000 |
| Oligohydramnios | 22 (2.579%) | 0 (0.0%) | 0.074 |
| **Placental Pathology** |  |  |  |
| Maternal vascular malperfusion |  |  |  |
| Vasculopathy |  |  | 0.000 |
| Classic (Macrophage) | 4 (0.522%) | 5 (3.106%) | 0.009 |
| Classic (Trophoblast) | 177 (23.289%) | 42 (26.087%) | 0.512 |
| Mixed type | 29 (3.816%) | 15 (9.317%) | 0.006 |
| Mural hypertrophy | 54 (7.105%) | 14 (8.696%) | 0.593 |
| Placental weight (grams) | 461.000 [391.000;531.000] | 464.000 [375.000;534.500] | 0.755 |
| Infarcts | 47 (5.510%) | 24 (14.634%) | 0.000 |
| Thrombosis | 196 (22.978%) | 42 (25.767%) | 0.503 |
| Subchorionic | 80 (9.379%) | 12 (7.317%) | 0.487 |
| Abruption | 13 (1.524%) | 3 (1.829%) | 1.000 |
| Fetal vascular malperfusion |  |  |  |
| Avascular villi | 113 (13.247%) | 16 (9.756%) | 0.270 |
| Inflammatory/infectious |  |  |  |
| MIR (acute chorioamnionitis) | 266 (31.184%) | 28 (17.073%) | 0.000 |
| MIR (chronic deciduitis) | 206 (24.150%) | 22 (13.415%) | 0.004 |
| FIR (acute funisitis/vasculitis) | 109 (12.778%) | 11 (6.707%) | 0.038 |
| Chronic villitis | 175 (20.516%) | 15 (9.146%) | 0.001 |
| **Other placental pathology** |  |  |  |
| Meconium | 249 (29.225%) | 20 (12.195%) | 0.000 |
| MFI/MPFD | 25 (2.934%) | 5 (3.049%) | 1.000 |
| **Lab tests and others** |  |  |  |
| Total WBC counts | 9.900 [8.300;12.300] | 9.050 [7.600;10.600] | 0.000 |
| Neutrophil differential (%) | 72.573 ± 7.363 | 70.679 ± 7.126 | 0.004 |
| Lymphocyte differential (%) | 16.950 [12.650;21.200] | 19.150 [14.900;23.400] | 0.000 |
| Body temperature (°C) | 36.700 [36.500;36.900] | 36.700 [36.600;37.000] | 0.129 |
| BMI | 30.385 [27.262;34.574] | 33.752 [29.485;37.637] | 0.000 |
| Blood pressure (Systolic) | 124.000 [117.000;132.000] | 142.000 [135.000;148.500] | 0.000 |
| Blood pressure (Diastolic) | 76.000 [70.000;82.000] | 89.000 [84.000;96.000] | 0.000 |

Separately, we have collected 619 placentas of late onset preeclampsia and 133 normal controls to determine the placental weight and associated various type of decidual vasculopathy [11]. The data grouped the classic type vasculopathy as one entity, and did not separate atherosis of macrophage type from atherosis of trophoblast type (Figure 4). There is a statistically significant association between classic type vasculopathy and the placental weight. Classic type vasculopathy is associated with lower placental weight in comparison to normal healthy pregnant controls. Mixed type vasculopathy is statistically associated with the lowest placental weight.

**Figure 4.**
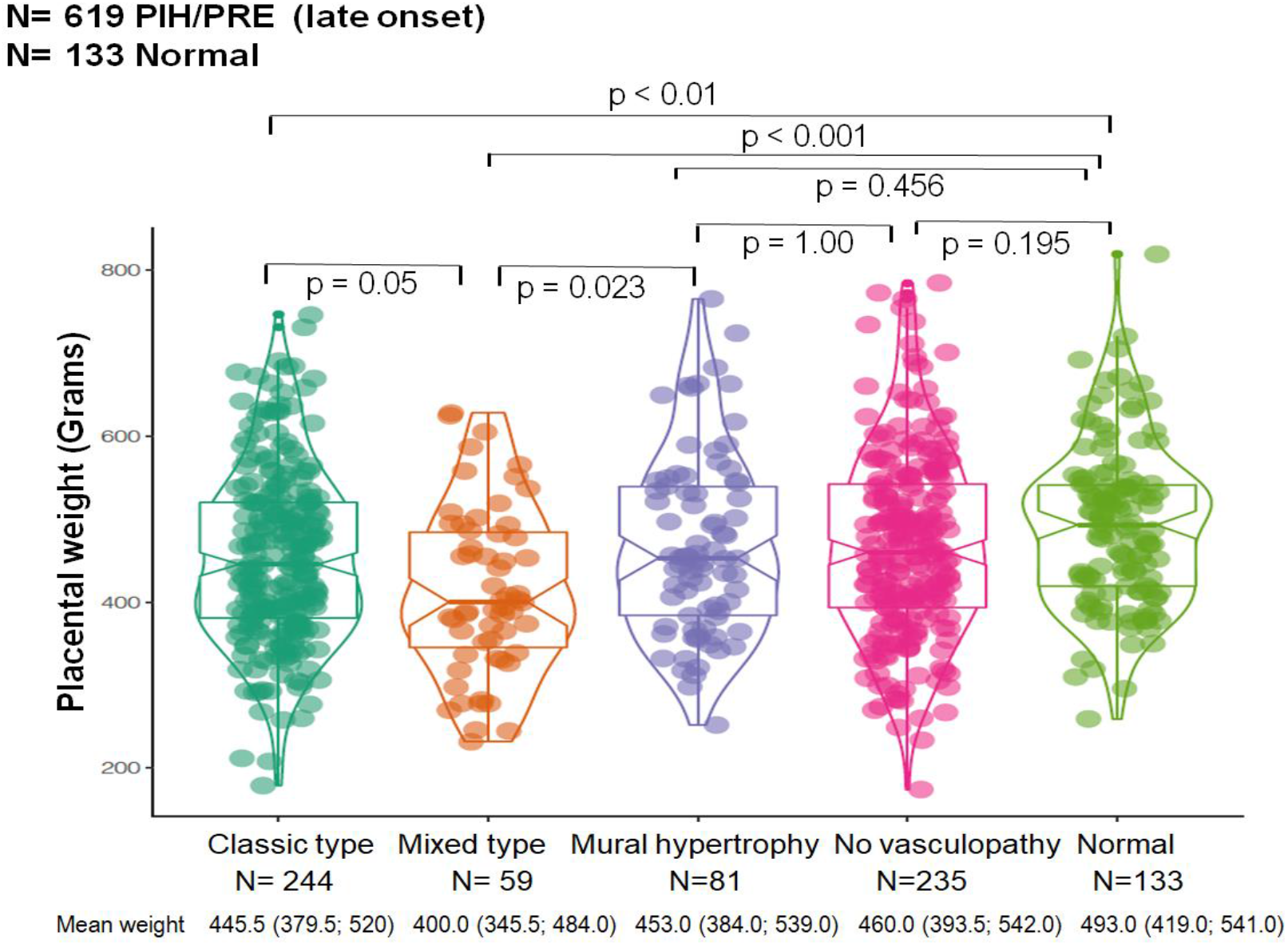
Placental weight versus various types of decidual vasculopathy in placentas of late onset preeclampsia in comparison to normal term placental controls. Statistical analysis was performed by using multi-variant ANOVA test in R-package. Mean placental weight and numbers in brackets represent 95% confidence interval (CI). P<0.05 represents statistically significant difference shown on the top of the chart.

## Discussion

Acute atherosis has been first described by Zeek in 1950, and there is very little debate about the diagnostic criteria or clinical significance of Zeek’s atherosis [6]. Acute atherosis has been associated with preeclampsia or hypertensive disorders of pregnancy, and numerous subsequent studies have confirmed this association, although the pathogenesis is unclear [6, 9, 10, 18, 19]. Oxidative stress of placental bed vessels in preeclampsia has been extensively studied due to the similarity of acute atherosis with adult atherosclerosis and it is clear that oxidative stress plays important roles in formation of atherosis (macrophage migration and deposit within the arterial wall) [8, 20–22]. It is also clear that lipid plays a role in formation of foamy macrophages in atherosis (oil-red-O), similar to atherosclerosis [23–25].

Atherosis of trophoblast type is so named that in a subset of acute atherosis previously described, the intramural or endovascular foamy cells are of trophoblast origin, rather than macrophage origin [11, 19, 26]. Identification of some of the foamy cells to be trophoblast origin leads to a different set of questions of pathogenesis. Endovascular or intramural trophoblasts have been described many decades ago, and identification of these trophoblasts under light microscope presents no problem for pathologists [5]. The critical link between the endovascular or intramural trophoblasts to foamy cells and atherosis is that vascular trophoblasts express CD56 exclusively, in addition to conventional trophoblast markers such as cytokeratins, p63 and HLA-G, and these vascular trophoblasts are originated from the “plugging cells” of spiral artery in early pregnancy [11, 27]. Spiral artery remodeling is an essential physiological step of normal pregnancy. Naturally, the presence of vascular trophoblasts in late pregnancy has been used as evidence of “remodeled spiral artery” in contrast to failed spiral artery remodeling. Remodeled spiral artery in late pregnancy has been called “physiologic change”[9, 15, 16, 19]. However, plugging cells from early implantation are known to disappear from approximately 7 weeks and the uteroplacental circulation is fully established at the end of first trimester [27]. Vascular trophoblasts are rarely seen in the third trimester of normal pregnancy [5, 11]. Temporal and spatial cellular functions of vascular trophoblasts have not been taken into consideration in the historic studies [15, 28]. Persistence of vascular trophoblasts in late pregnancy (third trimester) therefore represents abnormal (pathological) changes in placenta rather than normal physiologic features. However, persistence of vascular trophoblasts in late pregnancy has not been systemically studied to date due to reasons to our knowledge.

For a pathologist, the diagnostic criterion is critical for accurately categorizing the placental abnormalities. The changes in atherosis of trophoblast type should be similar to those of atherosis of macrophage type, ie, foamy cells in the background of fibrinoid medial necrosis of the arterial wall. In some cases, vascular trophoblasts are present without fibrinoid medial necrosis. These morphologic changes should also be called atherosis of trophoblast type, given the pathogenic mechanism remains as persistence of vascular trophoblasts in late pregnancy.

As atherosis of trophoblast type represents persistence of vascular trophoblasts in maternal circulation, we have attempted to examine the role of sex dimorphism in our data. Our current data show no evidence of sex difference in frequencies of atherosis of trophoblast type. However, study of sex dimorphism requires a large number of cases to achieve meaningful difference [29–31]. Our current dataset may not contain enough cases, and the data collection is still on-going with a potential update once we have adequate case number.

Our current data indicate that atherosis of trophoblast type is abnormal finding in placental examination, and should be stated as such in pathology report. Persistence of vascular trophoblasts invariably affects the vascular involution after delivery, and it has been associated with post-partum hemorrhage [32]. It is unclear how long these vascular trophoblasts persist in maternal arterial wall after delivery/pregnancy, and the presence of these vascular trophoblasts within the maternal spiral artery wall will likely affect the vascular stiffness and hemodynamics of maternal circulation after pregnancy [33–35]. The constellation of these maternal vascular changes is likely a significant risk factor for cardiovascular disease in later life after pregnancy. Further evidence is required to better assess the potential link of these morphologic changes of placental examination to better health of women after pregnancy.

## Conclusion

Atherosis of trophoblast type represents a spectrum of morphologic changes consistent with abnormal placental pathology, and it is associated with various maternal pregnancy complications with distinct pathogenic mechanism related to trophoblast cell death and vascular regeneration.

